# PERFECTO: Prediction of Extended Response and Growth Functions for Estimating Chemotherapy Outcomes in Breast Cancer

**DOI:** 10.1101/2020.12.29.424759

**Authors:** Daria Kurz, Cristian Axenie

**Affiliations:** Interdisciplinary Breast Center, Helios Klinikum Munich West, Munich, Germany; Audi Konfuzius-Institut Ingolstadt Lab, Technische Hochschule Ingolstadt Ingolstadt, Germany

**Keywords:** Machine Learning, Tumor Growth, Cancer Therapy Outcome, Breast Cancer, Personalized Medicine, Mathematical Oncology

## Abstract

Understanding tumor’s evolution under chemotherapy is central in the design of cancer therapy regimens. Drug resistance poses a major obstacle in the battle against most types of cancer and therapy design. Personalized treatments have the potential to offer greater effectiveness and the ability to prevent and circumvent drug resistance. In this study we introduce PERFECTO (Prediction of Extended Response and Growth Functions for Estimating ChemoTherapy Outcomes), a machine learning system capable of extracting the tumor growth function and response under chemotherapy. Exploiting the underlying correlations in the clinical data, the system captures the statistical peculiarities of tumor growth in-vivo without an explicit modeling of tumor microenvironment and expensive clinical investigations. We demonstrate the learning capabilities of PERFECTO in predicting unperturbed tumor growth and chemotherapy tumor growth from multiple clinical breast cancer datasets. We postulate that predictability is the key. Using PERFECTO clinicians will be able to improve treatment plans for patient-specific parameters from individual tumors. Our preliminary experiments on in-vitro, animal and in-vivo datasets, shown that, with a high degree of confidence, PERFECTO is able to estimate treatment effectiveness through an accurate tumor growth response prediction, independent of the breast cancer cell line. This in turn can alleviate the need of ordering extra clinical tests or any extra wait time before treatment initiation.

## I. Background

Chemotherapy regimens are chosen primarily based on empirical data from clinical trials and depend on patient’s form and subtype of cancer, progression to metastases, high-risk indications, and prognosis [1]. Such treatment plans consider consensus expert panel guidelines and laboratory-based testing that describe the variation in tumor tolerance and methods for the use of chemotoxic agents depending on their clinical efficacy [2]. Yet, physicians still face major challenges in successfully predicting the effectiveness (i.e. size of the tumor after neoadjuvant chemotherapy) of any particular chemotherapy plan for any given patient before the initiation of the regimen [3].

### A. Chemotherapy regimen planning: in-vitro vs. in-vivo prediction

In designing chemotherapy regimens, a critical problem faced by oncologists is the discrepancy between experiments performed in the laboratory on monolayers of cancer cells and treatments performed in live patients or on animals [4]. Various cytostatics exhibit highly effective results when delivered to controlled monolayers, but they under perform in animal models [5]. In theory, if the chemotherapeutic drug can reach and act upon the tumor in-vivo in the same concentrations and duration as in monolayer experiments, the amount of tumor cell death as a result of treatment should be comparable to that of the monolayer experiments.

But in-vivo cancers are resistant to systemic chemotherapy. The first reason is that, as tumors advance, their growth fraction decreases. Hence, as chemotherapy preferentially targets rapidly dividing cells, treatment resistance builds up [6]. Second, resistance to basically any drug can be attained through random genetic mutations accompanied by natural selection, resulting in tumors that are readily resistant to treatment [7]. Finally, physical properties of a tumor’s microenvironment (e.g. acidosis, hypoxia, tissue density, high interstitial fluid pressure, drug wash-out, or impaired blood flow) influence a drug’s ability to penetrate and kill tumor cells [8]. Such properties can be potential obstructions to drug diffusion, which increase the tumor’s resistance to chemotherapy [9]. Estimating the impact of such properties on tumor growth invivo is crucial for predicting therapy outcome.

Despite their importance, measuring such parameters invivo is not feasible [10] and mathematical models were developed in an attempt to capture their impact on tumor growth [11], [12]. In this context, the need for multimodal tumor growth data timeseries [13], expensive investigations (e.g. MRI, genetics) generating only days-level granularity timeseries [14], unevenly sampled measurements [15], variability in administered drug effect type on tumor growth [16], yield integrative computational models to support large-scale adoption in clinical use [17].

### B. Models of tumor growth

Various models that capture unperturbed tumor growth (i.e. not under cytostatic administration) in-vitro and in live animals have been developed [18]–[20]. Capturing the biological substrate of physical processes in cancer cells, such models have dominated the computational oncology field. Ordinary Differential Equations (ODE) tumor growth models [21] are typically the norm. In our study, we explored three of the most representative and typically used scalar growth models, namely Logistic, von Bertalanffy, and Gompertz, described in Table I-B. Such models are typically used to predict the kinetics of unperturbed tumors. Studies of treatments using such models demonstrated that such growth equations cannot be directly reapplied to determine tumor growth trajectory [23]. Nor can they, then, necessarily be used to predict chemotherapy outcome, as treatment fundamentally alters the growth trajectory by changing the asymptotic tumor size [24].

**TABLE I:** Overview of tumor growth models *f* (*V*) in our study. Parameters: *V* - volume (or cell population size through conversion - 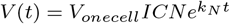 where *N* is the population size, *ICN* is the initial cell number, *V*_*onecell*_ is the volume of one cell and *k*_*N*_ is the rate constant for changes in cell number as considered in [22]), *α* - growth rate, *β* - cell death rate, *λ* - nutrient limited proliferation rate, *k* - carrying capacity of cells.

| Model | Equation |
| --- | --- |
| Logistic [18] | $f_L(V) = \frac{dV}{dt} = \alpha V - \beta V^2$ |
| Bertalanffy [19] | $f_B(V) = \frac{dV}{dt} = \alpha V^\lambda - \beta V$ |
| Gompertz [20] | $f_G(V) = \frac{dV}{dt} = V(\beta - \alpha \ln V)$ |

### C. Objectives of the study

The objective of our study is to explore the capabilities of a machine learning model to capture tumor growth in both unperturbed and under neoadjuvant chemotherapy. By employing only tumor growth timeseries data (i.e. tumor volume measurements from different sources: MRI, caliper, ultrasound), the model should learn the tumor growth function and tumor’s extended response during chemotherapy. Hence, capturing the effect cytostatics have upon the tumor.

Using publicly available clinical datasets for tumor growth in-vitro, animal xenografts, and humans, we demonstrate that PERFECTO is able to:

- learn the unperturbed tumor growth function from tumor growth data of different breast cancer cell lines;
- learn the tumor growth function and response under chemotherapy regimen in humans; and
- predict chemotherapy outcome more precisely than traditional growth models in a data-driven manner, without biologically accurate parametrization.

## II. Materials and methods

In the next section we introduce the underlying mechanisms of PERFECTO as well as the procedures used in our experiments.

### A. Introducing PERFECTO

PERFECTO is an unsupervised machine learning system composed of an encoder and a correlation detector. The encoder, is responsible to map the input data timeseries to a latent space in order to capture and preserve the statistics of the input space. The correlation detector is responsible of acting upon multiple such (encoded) latent spaces and extract their temporal correlation. At design time, one can use, for instance, Gaussian process prior variational autoencoders [25] or Correlated Variational Auto-Encoders (CVAEs) [26]. In the current instantiation of PERFECTO, for simplicity, we consider a Self-Organizing Maps (SOM)-based encoder [27] and a Hebbian Learning (HL) [28] correlation detector. We introduce the basic functionality of PERFECTO through a simple example in Figure 1. Here, we use measurements from 49 histology images of metastatic cancer to liver in the cohort of patients from [11]. The two input timeseries (i.e. direct measurement of fraction of tumor kill and kill radius from histopathology of biopsies after chemotherapy) follow a second order (i.e. parabolic) dependency, depicted in Figure 1a. As depicted in Figure 1, PERFECTO is able to learn the extended response of the tumor to chemotherapy from timeseries data of fraction of tumor kill and kill radius from histopathology of biopsies.

**Fig. 1.**
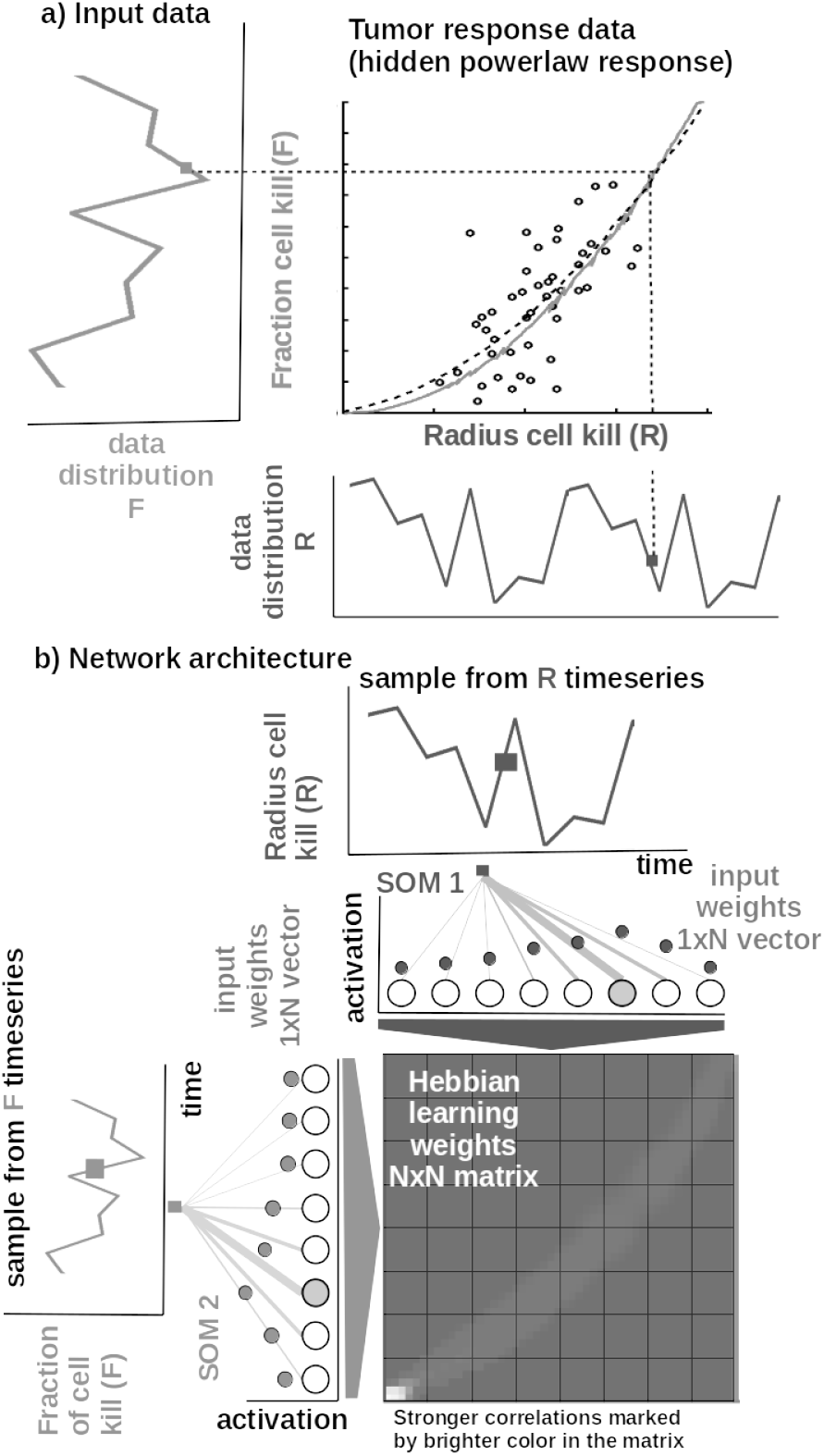
Basic functionality of PERFECTO: a) Tumor response data resembling a non-linear relation (i.e. parabola) hidden in the timeseries (i.e. fraction of cell kill (F) vs. radius of cell kill (R)) learnt by PERFECTO (plain trace) w.r.t patient data (circles) and model fit (dashed line) from [11]. b) Basic architecture of PERFECTO: 1D SOM encoder networks with *N* neurons encoding the timeseries (i.e. fraction of tumor kill vs. kill radius), and a *NxN* Hebbian connection matrix coupling the two 1D SOMs that will eventually encode the relation between the timeseries, i.e. extended response.

#### Core model

The input SOMs (i.e. 1D lattice networks with *N* neurons) encode timeseries samples in a distributed activity pattern, as shown in Figure 1b. This activity pattern is generated such that the closest preferred value of a neuron to the input sample will be strongly activated and will decay, proportional with distance, for neighbouring units. The SOM specializes to represent a certain (preferred) value in the timeseries and learns its sensitivity, by updating its tuning curves shape. Given an input sample *s^p^*(*k*) from one timeseries at time step *k*, the network computes for each *i*-th neuron in the *p*-th input SOM (with preferred value 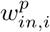 and tuning curve size 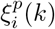 the elicited neural activation as

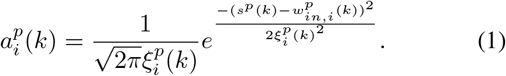

The winning neuron of the *p*-th population, *b^p^*(*k*), is the one which elicits the highest activation given the timeseries sample at time *k*

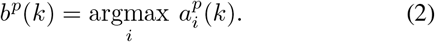

The competition for highest activation in the SOM encoder is followed by cooperation in representing the input space. Hence, given the winning neuron, *b^p^*(*k*), the cooperation kernel,

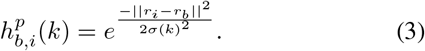

allows neighbouring neurons (i.e. found at position *r_i_* in the network) to precisely represent the input sample given their location in the neighbourhood *σ*(*k*) of the winning neuron. The neighbourhood width *σ*(*k*) decays in time, to avoid twisting effects in the SOM. The cooperation kernel in Equation 3, ensures that specific neurons in the network specialize on different areas in the input space, such that the input weights (i.e. preferred values) of the neurons are pulled closer to the input sample,

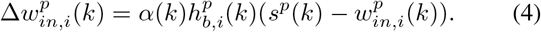

Neurons in the two encoder SOMs are then linked by a fully (all-to-all) connected matrix of synaptic connections, where the weights in the matrix are computed using Hebbian learning. The connections between uncorrelated (or weakly correlated) neurons in each population (i.e. *w_cross_*) are suppressed (i.e. darker color) while correlated neurons connections are enhanced (i.e. brighter color), as depicted in Figure 1b. Formally, the connection weight 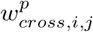 between neurons *i, j* in the different input SOMs are updated with a Hebbian learning rule as follows:

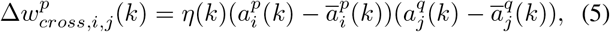

where 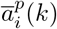 is a “momentum” like exponential moving average. Hebbian learning ensures that when neurons fire synchronously their connection strengths increase, whereas if their firing patterns are anti-correlated the weights decrease. The weight matrix encodes the co-activation patterns between the input layers (i.e. SOMs), as shown in Figure 1b, and, eventually, the learned growth law (i.e. functional relation) given the timeseries, as shown in Figure 1a. Self-organisation and Hebbian correlation learning processes evolve simultaneously, such that both the representation and the extracted relation are continuously refined, as new samples are presented. This can be observed in the encoding and decoding functions where the input activations are projected though *w_in_* (Equation 1) to the Hebbian matrix and then decoded through *w_cross_*.

#### Parametrization and read-out

In all of our experiments the data is fed to PERFECTO which encodes each timeseries in the SOM encoders and learns the underlying relation using the Hebbian correlation detector. In our experiments, each of the SOM has *N* = 50 neurons, the Hebbian connection matrix has size *NxN* and parametrization is done as: *alpha* = [0.01, 0.1] decaying, *η* = 0.9, 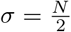 decaying following an inverse time law. We use as decoding mechanism an optimization method that recovers the real-world value given the self-calculated bounds of the input timeseries. The bounds are obtained as minimum and maximum of a cost function of the distance between the current preferred value of the winning neuron (i.e. the value in the input which is closest to the weight vector of the neuron in Euclidean distance) and the input sample.

### B. Datasets

In our initial experiments we used publicly available tumor growth datasets from in-vitro and animal xenografts (see Table II), with real clinical tumor volume measurements, for different cell lines of breast cancer. This choice is to probe and demonstrate the versatility of PERFECTO in learning from tumor growth patterns induced by different types of cancer. In the last series of experiments, we used real patient data from the I-SPY 1 TRIAL: ACRIN 6657 [32]. Data for the 136 patients treated for breast cancer in the IPSY-1 clinical trial was obtained from the cancer imaging archive ^1^ and the Breast Imaging Research Program at UCSF. The timeseries data contained only the largest tumor volume from MRI measured before therapy, 1 to 3 days after therapy, between therapy cycles, and before surgery, respectively. To summarize, the properties of the dataset are depicted in Figure 2.

**TABLE II:** Description of the breast cancer datasets used in the experiments.

| Experimental dataset setup |  |  |  |  |  |
| --- | --- | --- | --- | --- | --- |
| Dataset | Cancer Type | Data Type | Data Points | Data Freq. | Source |
| 1 | MDA-MB-231 cell line | Fluorescence imaging | 7 | 2x/week | [29] |
| 2 | MDA-MB-435 cell line | Digital Caliper | 14 | 2x/week | [15] |
| 3 | MCF-7 cell line | Caliper | 8 | 1x/week | [30] |
| 4 | LM2-4LUC+ cell line | Digital Caliper | 10 | 3x/week | [31] |

**Fig. 2.**
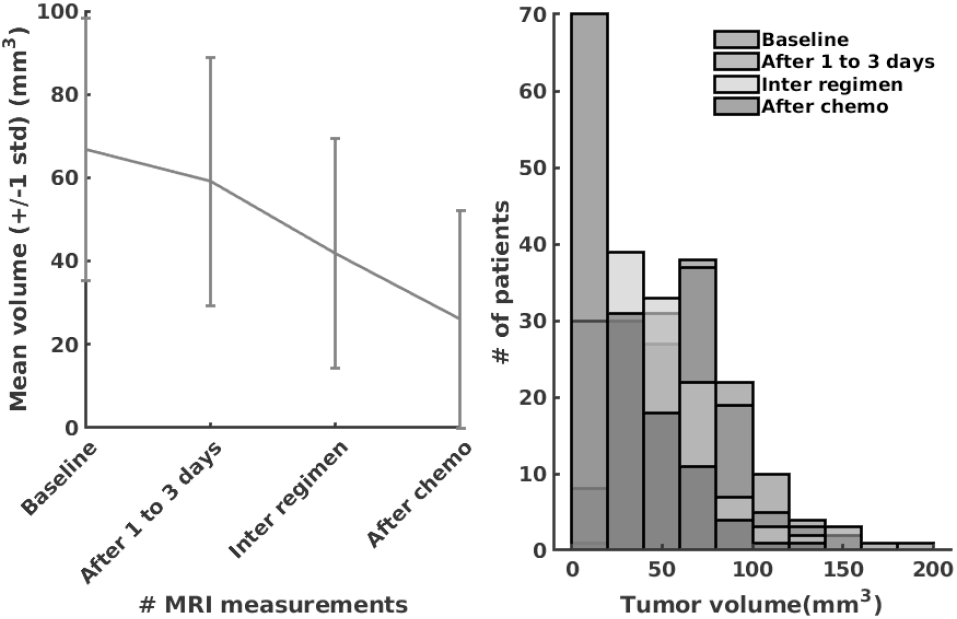
The I-SPY2 Trial Dataset properties.

### C. Procedures

In order to reproduce the experiments in our study, the MATLAB ^®^ code and copies of all the datasets are available on GITLAB ^2^. Each of the three mechanistic tumor growth models (i.e. Logistic, Bertalanffy, Gompertz) and PERFECTO were presented the tumor growth data in each of the clinical datasets. When a dataset contained multiple trials and patients, training was performed on 80% of records and 20% of records were used for testing and validation.

#### Mechanistic models setup

Each of the three ODE tumor growth models was implemented as ordinary differential equation (ODE) and integrated over the dataset length. We used a solver based on a modified Rosenbrock formula of order 2 that evaluates the Jacobian during each step of the integration. To provide initial values and the best parameters (i.e. *α, β, λ, k*) for each of the four models the Nelder-Mead simplex direct search (i.e. derivative-free minimum of unconstrained multi-variable functions) was used, with a termination tolerance of 10*e*^−4^ and upper bounded to 1000 iterations. Finally, fitting was performed by minimizing the sum of squared residuals.

#### PERFECTO setup

For PERFECTO the data was normalized before training and de-normalized for the evaluation. The system was comprised of two input SOM encoders, each with *N* = 50 neurons, encoding the volume data and the irregular sampling time sequence, respectively. Both input density learning and correlation learning cycles were bound to 100 epochs.

## III. Results

In the current section, we present the experimental results of our study and demonstrate that PERFECTO is capable to: a) learn the unperturbed tumor growth function from tumor growth data of various cell lines of breast cancer, b) learn the tumor growth function and response under chemotherapy, and c) to use the learnt quantities to provide a precise outcome of the therapy (i.e. accurate tumor size before surgery).

### A. Learning the unperturbed tumor growth function

The first experiments addressed the capability to learn the tumor growth function *f* (*V*) from data without imposing biological constraints upon the cell line, tumor size, number of cells etc. We demonstrate the superior learning capabilities of PERFECTO on the four publicly-available breast cancer clinical datasets described in Table II. As we can observe in Table III, PERFECTO learns a superior fit (i.e. lowest Sum Squared Error (SSE), Root Mean Squared Error (RMSE) and symmetric Mean Absolute Percentage Error (sMAPE)) to the tumor growth data, despite the limited number of samples (i.e. 7 data points for MDA-MD-231 cell line dataset and up to 14 data points for MDA-MD-435 cell line dataset). Despite their ubiquitous use, the classical tumor growth models (e.g. Gompertz, von Bertalanffy, Logistic) are confined due to: a) the requirement of a precise biological description - as one can see in the different sigmoid shapes in Figure 3; b) incapacity to describe the diversity of tumor types, and; c) the small amount and irregular sampling of the data - visible in the relatively poor fit to the data, captured in Figure 3.

**TABLE III:** Evaluation of tumor growth models in breast cancer.

|  |  | Evaluation Metrics |  |  |
| --- | --- | --- | --- | --- |
|  | Dataset/Model | SSE | RMSE | sMAPE |
| *MDA-MB-231 cell line cancer [29]* |  |  |  |  |
|  | Logistic | 7009.6 | 37.4423 | 1.7088 |
|  | Bertalanffy | 8004.9 | 44.7350 | 1.7088 |
|  | Gompertz | 7971.8 | 39.9294 | 1.7088 |
|  | **PERFECTO** | 119.3 | 4.1285 | 0.0768 |
| *MDA-MB-435 cell line cancer [15]* |  |  |  |  |
|  | Logistic | 0.2936 | 0.1713 | 0.1437 |
|  | Bertalanffy | 0.2315 | 0.1604 | 0.1437 |
|  | Gompertz | 0.3175 | 0.1782 | 0.1437 |
|  | **PERFECTO** | 0.0977 | 0.0902 | 0.0763 |
| *MCF-7 cell line cancer [30]* |  |  |  |  |
|  | Logistic | 3.0007 | 0.7072 | 1.0607 |
|  | Bertalanffy | 3.2943 | 0.8117 | 1.0607 |
|  | Gompertz | 3.1909 | 0.7293 | 1.0607 |
|  | **PERFECTO** | 0.7669 | 0.3096 | 0.2615 |
| *LM2-4LUC+ cell line cancer [31]* |  |  |  |  |
|  | Logistic | 45.6032 | 2.3876 | 1.4816 |
|  | Bertalanffy | 56.0739 | 2.8303 | 1.4816 |
|  | Gompertz | 53.2428 | 2.5798 | 1.4816 |
|  | **PERFECTO** | 0.2009 | 0.1417 | 0.0365 |
| *I-SPY2 Trial [32]* |  |  |  |  |
|  | Logistic | 248.3735 | 11.1439 | 1.7833 |
|  | Bertalanffy | 259.0963 | 16.0963 | 1.7834 |
|  | Gompertz | 260.3747 | 11.4100 | 1.7883 |
|  | **PERFECTO** | 0.8650 | 0.4650 | 0.0389 |

**Fig. 3.**
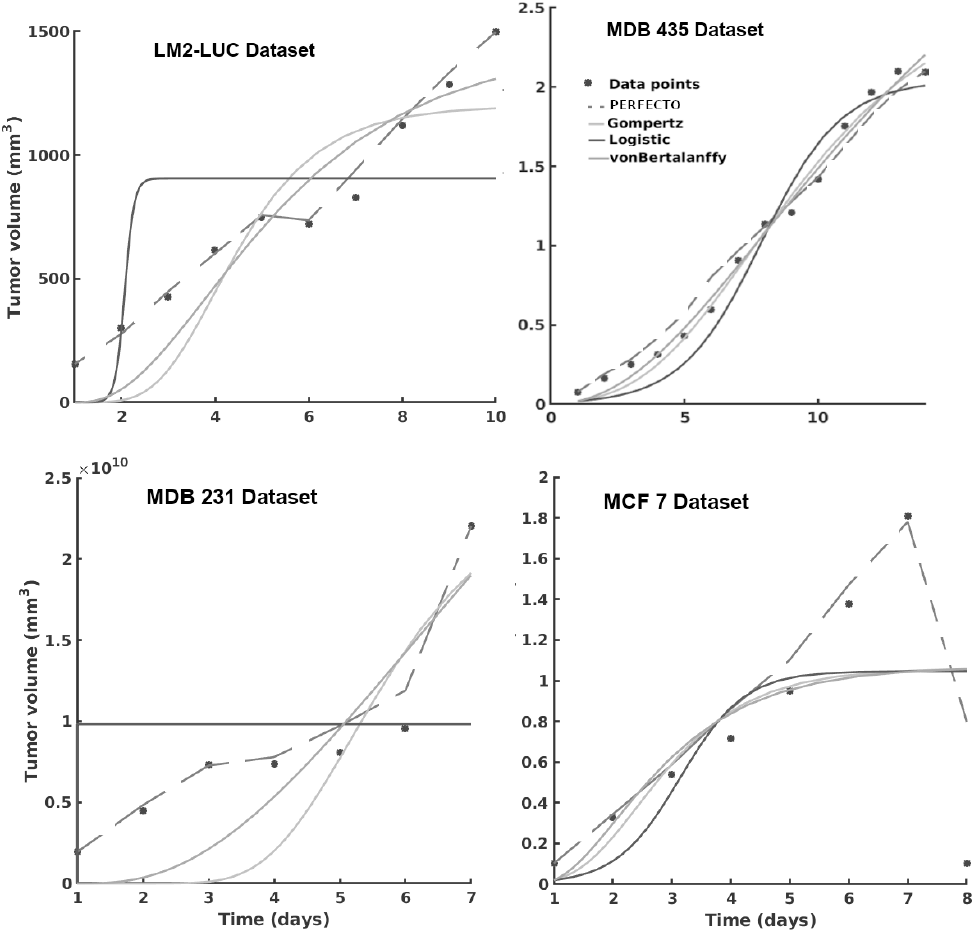
Evaluation of the tumor growth models on the different datasets: learnt growth functions. PERFECTO is decoded from the learnt Hebbian weight matrix. The decrease in the MCF7 Dataset is due to the administered chemotherapy and demonstrates the adaptivity of PERFECTO in capturing growth behaviors.

### B. Learning the tumor growth function and response under chemotherapy

When analyzing tumor growth functions and response under chemotherapy, we faced the high variability among patients given by the typical constellation of hormone receptors indicators (i.e. HR and HER2neu, which covered the full spectrum of positive and negative values) for positive and negative prognosis.

As we can observe in Table III, PERFECTO learns a superior fit overall the three metrics, capturing the intrinsic impact chemotherapy has upon the tumor growth function, despite the limited number of samples (i.e. 4 data points of the dataset overall evaluation dataset of 20% of patients).

## IV. Conclusion

Oncologists predict which drugs, concentrations, and delivery times will work best based on guidelines generated from patient statistics, animal experiments, or cell culture experiments. Yet, predicting how a tumor will react to a certain therapy regimen is still resistant to straightforward solutions. Either extrapolating from in-vitro pharmacokinetics models, to modeling tumor microenvironments, predicting tumor growth, in both unperturbed and under therapy regimens, poses challenges. In this study, we introduced and evaluated a machine learning model capable of predicting tumor growth, in both unperturbed context and under cytostatics, that outperforms traditional mechanistic models on real-world linical datasets. Such a solution has the potential to alleviate the unnecessary side effects from excessive dosages and to shape subsequent cycles or therapy decisions specific to a certain patient.

## Footnotes

1 https://wiki.cancerimagingarchive.net/display/Public/ISPY1

2 https://gitlab.com/akii-microlab/perfecto

## References

[1] F. Cardoso, E. Senkus, A. Costa, E. Papadopoulos, M. Aapro, F. André, N. Harbeck, B. Aguilar Lopez, C. Barrios, J. Bergh et al., “4th eso–esmo international consensus guidelines for advanced breast cancer (abc 4),” Annals of Oncology, vol. 29, no. 8, pp. 1634–1657, 2018.

[2] F. Cardoso, S. Kyriakides, S. Ohno, F. Penault-Llorca, P. Poortmans, Rubio, S. Zackrisson, and E. Senkus, “Early breast cancer: Esmo clinical practice guidelines for diagnosis, treatment and follow-up,” Annals of Oncology, vol. 30, no. 8, pp. 1194–1220, 2019.

[3] L. J. Oostendorp, P. F. Stalmeier, A. R. T. Donders, W. T. van der Graaf, and P. B. Ottevanger, “Efficacy and safety of palliative chemotherapy for patients with advanced breast cancer pretreated with anthracyclines and taxanes: a systematic review,” The Lancet Oncology, vol. 12, no. 11, pp. 1053–1061, 2011.

[4] C. Pauli, B. D. Hopkins, D. Prandi, R. Shaw, T. Fedrizzi, A. Sboner, Sailer, M. Augello, L. Puca, R. Rosati et al., “Personalized in vitro and in vivo cancer models to guide precision medicine,” Cancer discovery, vol. 7, no. 5, pp. 462–477, 2017.

[5] Y. Wu, Z. Deng, H. Wang, W. Ma, C. Zhou, and S. Zhang, “Repeated cycles of 5-fluorouracil chemotherapy impaired anti-tumor functions of cytotoxic t cells in a ct26 tumor-bearing mouse model,” BMC immunology, vol. 17, no. 1, p. 29, 2016.

[6] D. A. Kessler, R. H. Austin, and H. Levine, “Resistance to chemotherapy: patient variability and cellular heterogeneity,” Cancer research, vol. 74, no. 17, pp. 4663–4670, 2014.

[7] N. E. Navin, “Tumor evolution in response to chemotherapy: phenotype versus genotype,” Cell reports, vol. 6, no. 3, pp. 417–419, 2014.

[8] E. Henke, R. Nandigama, and S. Ergün, “Extracellular matrix in the tumor microenvironment and its impact on cancer therapy,” Frontiers in Molecular Biosciences, vol. 6, p. 160, 2020.

[9] J. K. Saggar, M. Yu, Q. Tan, and I. F. Tannock, “The tumor microenvironment and strategies to improve drug distribution,” Frontiers in oncology, vol. 3, p. 154, 2013.

[10] A. I. Minchinton and I. F. Tannock, “Drug penetration in solid tumours,” Nature Reviews Cancer, vol. 6, no. 8, pp. 583–592, 2006.

[11] J. Pascal, E. L. Bearer, Z. Wang, E. J. Koay, S. A. Curley, and V. Cristini, “Mechanistic patient-specific predictive correlation of tumor drug response with microenvironment and perfusion measurements,” Proceedings of the National Academy of Sciences, vol. 110, no. 35, pp. 14 266–14 271, 2013.

[12] Y. Cui, X.-P. Zhang, Y.-S. Sun, L. Tang, and L. Shen, “Apparent diffusion coefficient: potential imaging biomarker for prediction and early detection of response to chemotherapy in hepatic metastases,” Radiology, vol. 248, no. 3, pp. 894–900, 2008.

[13] D. Abler, P. Büchler, and R. C. Rockne, “Towards model-based characterization of biomechanical tumor growth phenotypes,” in Mathematical and Computational Oncology, G. Bebis, T. Benos, K. Chen, K. Jahn, and E. Lima, Eds. Cham: Springer International Publishing, 2019, pp. 75–86.

[14] C. L. Roland, S. P. Dineen, K. D. Lynn, L. A. Sullivan, M. T. Dellinger, L. Sadegh, J. P. Sullivan, D. S. Shames, and R. A. Brekken, “Inhibition of vascular endothelial growth factor reduces angiogenesis and modulates immune cell infiltration of orthotopic breast cancer xenografts,” Molecular Cancer Therapeutics, vol. 8, no. 7, pp. 1761–1771, 2009.

[15] L. D. Volk, M. J. Flister, D. Chihade, N. Desai, V. Trieu, and S. Ran, “Synergy of nab-paclitaxel and bevacizumab in eradicating large orthotopic breast tumors and preexisting metastases,” Neoplasia, vol. 13, no. 4, pp. 327–IN14, 2011.

[16] T. D. Gaddy, Q. Wu, A. D. Arnheim, and S. D. Finley, “Mechanistic modeling quantifies the influence of tumor growth kinetics on the response to anti-angiogenic treatment,” PLoS computational biology, vol. 13, no. 12, 2017.

[17] H. B. Frieboes, M. E. Edgerton, J. P. Fruehauf, F. R. Rose, L. K. Worrall, R. A. Gatenby, M. Ferrari, and V. Cristini, “Prediction of drug response in breast cancer using integrative experimental/computational modeling,” Cancer research, vol. 69, no. 10, pp. 4484–4492, 2009.

[18] P.-F. Verhulst, “Notice sur la loi que la population suit dans son accroissement,” Corresp. Math. Phys., vol. 10, pp. 113–126, 1838.

[19] L. Von Bertalanffy, “Quantitative laws in metabolism and growth,” The quarterly review of biology, vol. 32, no. 3, pp. 217–231, 1957.

[20] B. Gompertz, “On the nature of the function expressive of the law of human mortality, and on a new mode of determining the value of life contingencies. in a letter to francis baily, esq. frs &c,” Philosophical transactions of the Royal Society of London, no. 115, 1825.

[21] P. Gerlee, “The model muddle: in search of tumor growth laws,” Cancer research, vol. 73, no. 8, pp. 2407–2411, 2013.

[22] H.-J. Kuh et al., “Computational model of intracellular pharmacokinetics of paclitaxel,” Journal of Pharmacology and Experimental Therapeutics, vol. 293, no. 3, 2000.

[23] F. Kozusko and Ž. Bajzer, “Combining gompertzian growth and cell population dynamics,” Mathematical Biosciences, vol. 185, no. 2, pp. 153–167, 2003.

[24] F. Kozusko, M. Bourdeau, Z. Bajzer, and D. Dingli, “A microenvironment based model of antimitotic therapy of gompertzian tumor growth,” Bulletin of Mathematical Biology, vol. 69, no. 5, pp. 1691–1708, 2007.

[25] F. P. Casale, A. Dalca, L. Saglietti, J. Listgarten, and N. Fusi, “Gaussian process prior variational autoencoders,” in Advances in Neural Information Processing Systems, 2018, pp. 10 369–10 380.

[26] D. Tang, D. Liang, T. Jebara, and N. Ruozzi, “Correlated variational auto-encoders,” arXiv preprint arXiv:1905.05335, 2019.

[27] T. Kohonen, “Self-organized formation of topologically correct feature maps,” Biological cybernetics, vol. 43, no. 1, pp. 59–69, 1982.

[28] Z. Chen, S. Haykin, J. J. Eggermont, and S. Becker, Correlative learning: a basis for brain and adaptive systems. John Wiley & Sons, 2008.

[29] A. Rodallec, S. Giacometti, J. Ciccolini, and R. Fanciullino, “Tumor growth kinetics of human MDA-MB-231 cells transfected with dTomato lentivirus,” Dec. 2019. [Online]. Available: https://doi.org/10.5281/zenodo.3593919

[30] G. e. a. Tan, “Combination therapy of oncolytic herpes simplex virus hf10 and bevacizumab against experimental model of human breast carcinoma xenograft,” International Journal of Cancer, vol. 136, no. 7, pp. 1718–1730, 2015.

[31] M. Mastri, A. Tracz, and J. M. Ebos, “Tumor growth kinetics of human LM2-4LUC+ triple negative breast carcinoma cells,” Dec. 2019. [Online]. Available: https://doi.org/10.5281/zenodo.3574531

[32] D. Yee, A. M. DeMichele, C. Yau, C. Isaacs, W. F. Symmans, K. S. Albain, Y.-Y. Chen, G. Krings, S. Wei, S. Harada et al., “Association of event-free and distant recurrence–free survival with individual-level pathologic complete response in neoadjuvant treatment of stages 2 and 3 breast cancer: Three-year follow-up analysis for the i-spy2 adaptively randomized clinical trial,” JAMA oncology, 2020.

